# Mosquito aging modulates the development, virulence and transmission potential of pathogens

**DOI:** 10.1101/2023.09.21.558787

**Authors:** Bernard Somé, Edwige Guissou, Dari F. Da, Quentin Richard, Marc Choisy, Koudraogo Bienvenue Yameogo, Domombabele FdS Hien, Rakiswende S Yerbanga, Georges Anicet Ouedraogo, Kounbobr R Dabiré, Ramsès Djidjou-Demasse, Anna Cohuet, Thierry Lefèvre

## Abstract

Host age variation is a striking source of heterogeneity that can shape the evolution and transmission dynamic of pathogens. Compared to vertebrate systems, our understanding of the impact of host age on invertebrate-pathogen interactions remains limited. We examined the influence of mosquito age on key life-history traits driving human malaria transmission. Females of *Anopheles coluzzii*, a major malaria vector, belonging to three age classes (4, 8, and 12 day-old), were experimentally infected with *Plasmodium falciparum* field isolates. Our findings revealed reduced competence in 12-day-old mosquitoes, characterized by lower oocyst/sporozoite rates and intensities compared to younger mosquitoes. Despite shorter median longevities in older age classes, infected 12-day-old mosquitoes exhibited improved survival, suggesting that the infection might act as a fountain of youth for older mosquitoes specifically. The timing of sporozoite appearance in the salivary glands remained consistent across mosquito age classes, with an extrinsic incubation period of approximately 13 days. Integrating these results into an epidemiological model revealed a lower vectorial capacity for older mosquitoes compared to younger ones, albeit still substantial due to extended longevity in the presence of infection. Considering age heterogeneity provides valuable insights for ecological and epidemiological studies, informing targeted control strategies to mitigate pathogen transmission.

## Introduction

Hosts and parasites are engaged in ongoing coevolutionary interactions, which generates selection for resistant hosts and infectious parasites. Studies have shown that infection traits, such as infectivity, virulence, resistance and tolerance are genetically determined and their expression can depend on interactions between host and parasite genotypes [1–5]. However, the influence of parasite environment on infections traits is less well understood [6–8].

Age variation among hosts within a population is one of the most significant sources of heterogeneity, with profound implications for the expression and evolution of infection traits [9,10]. For instance, in human populations, age plays a critical role in determining host susceptibility and parasite virulence [11], particularly with pathogens causing prolonged and persistent symptoms or newly emerging pathogens [12]. In comparison to our understanding of age-specific patterns of infection in vertebrates, our understanding of the impact of host age on invertebrate-parasite interactions remains limited.

The relationship between host age and the outcome of infection in invertebrates is complex, with contrasting findings observed across various studies, organisms, and specific phenotypic traits. Despite the paucity of data available, recent research indicates that invertebrate hosts, unlike vertebrates, generally exhibit decreased susceptibility and infection-induced mortality with age [13]. Yet, the widespread occurrence of immunosenescence across the tree of life, including in insects [14], suggests the opposite; that is, a decline in immune function, and hence in parasite resistance with age [15,16]. In insects, studies have revealed that immunosenescence can manifest as excessive, overly-reactive and less specific immune responses, without any notable improvement in survival [17–19]. Furthermore, different components of the insect immune response (enzymatic versus cellular) can display distinct age-specific patterns [16].

Examining the impact of age on infection traits requires host organisms with age classes sharing the same habitat, consuming identical food resources and exhibiting overlapping generations, hence causing substantial age variation in their natural populations. Adult mosquitoes fulfill these criteria and are particularly appropriate to address the effects of host age on infection traits. Mosquitoes reach a definitive body size following emergence (i.e. no size variation with aging in adults), all age classes share the same terrestrial and aerial environment and feed on blood (for egg production) and plant-derived sugars (for energy and maintenance). With a rather short gonotrophic cycle, which can be as fast as 48h between two egg-lays, mosquito females are recurrently looking for a blood meal during which they can transmit pathogens responsible for devastating diseases, such as malaria, dengue fever or Zika [20].

In mosquitoes, as in most organisms, aging and the number of feeding episodes are inherently positively correlated, and both feeding- and age-related mechanisms can affect pathogen transmission. Firstly, the age structure of the mosquito population is a key determinant of the intensity of pathogen transmission [21–24]. Owing to the process of senescence, younger mosquitoes exhibit higher survival rates compared with older mosquitoes [25] and are therefore more likely to endure the relatively long extrinsic incubation period of pathogens (*i.e.*, the EIP, the time required to replicate and disseminate to the mosquito salivary glands after the ingestion of an infected bloodmeal). Although the EIP can vary widely in response to genetic and environmental factors, it typically ranges from 7 to 14 days for pathogens like *Plasmodium*, dengue and Zika [26–32]. This duration encompasses a considerable portion of a mosquito expected lifespan [33,34]. Secondly, mounting evidence points to the occurrence of immunosenescence in mosquitoes, characterized by declining levels of melanization [35–37] and circulating hemocytes [38,39] as individuals get older. These age-related declines are expected to coincide with increased infection levels in older mosquitoes [38], although conflicting observations have also been reported [39]. Thirdly, previous blood-feeding can effectively trigger innate immune responses and/or accelerate digestion of the infective blood meals in mosquitoes, resulting in decreased susceptibility to infection [40–43]. In natural conditions, where older mosquitoes have experienced a greater number of feeding episodes, it suggests a reduction in infection levels as mosquitoes age. Together these observations highlight the intricate interplay between mosquito aging, successive feeding cycles, and pathogen transmission.

Recent theoretical works emphasize the need to better understand and characterize the contribution of vector age structure to the heterogeneous transmission and virulence of vector-borne pathogens [23,44–46]. In order to gain insight on this matter, we examined whether key life-history traits that drive human malaria transmission vary with mosquito age. Vector competence for pathogens (i.e. a combined estimate of parasite infectivity and vector susceptibility), vector survival and the pathogen’s extrinsic incubation period (EIP) are three important components of vectorial capacity, a classic measure of the transmission potential of a vector-borne pathogen. The effect of vector age on each of these three traits was assessed using an epidemiologically-relevant vector-pathogen system consisting of the parasite *Plasmodium falciparum*, which causes the most severe form of human malaria, and the mosquito *Anopheles coluzzii*, a major vector of *P. falciparum* in Africa. Females of *Anopheles coluzzii* belonging to different age classes were infected with sympatric field isolates of *Plasmodium falciparum* using direct membrane feeding assays. Through a series of experiments, the effects of mosquito age on (i) mosquito competence, (ii) mosquito survival, (iii) and the parasite’s EIP were examined. These results were then combined into an epidemiological model to predict the contribution of mosquito age variation to overall malaria transmission.

## Materials and Methods

### Mosquitoes

Laboratory-reared *An. coluzzii* were obtained from an outbred colony established in 2008 and that have since been repeatedly replenished with wild-caught mosquito females collected in Soumousso, (11°23’14“N, 4°24’42”W), 40 km from Bobo Dioulasso, south-western Burkina Faso (West Africa), and identified by SINE PCR [47]. Mosquitoes were held in 30 cm × 30 cm × 30 cm mesh-covered cages and reared under standard insectary conditions (12:12 LD, 27 ± 2°C, 70 ± 5% relative humidity). Females were maintained on rabbit blood by direct feeding (protocol approved by the national committee of Burkina Faso; IRB registration #00004738 and FWA 00007038), and adult males and females fed with a 5% glucose solution. Larvae were reared at a density of about 300 first instar larvae in 700 ml of water in plastic trays and were fed with Tetramin Baby Fish Food (Tetrawerke, Melle, Germany).

### Parasite field isolates and mosquito experimental infection

*Anopheles coluzzii* females belonging to distinct age classes (see details below in each experiments) were fed with blood drawn from naturally *P. falciparum* gametocyte-infected patients recruited among 5–12-year-old school children in villages surrounding Bobo-Dioulasso, Burkina Faso, using Direct Membrane Feeding Assays (DMFA) as previously described [48–50]. Briefly, thick blood smears were taken from each volunteer, air-dried, giemsa-stained, and examined by microscopy for the presence of *P. falciparum*. Asexual trophozoite parasite stages were counted against 200 leucocytes, while infectious gametocytes stages were counted against 1000 leukocytes. Children with asexual parasitemia of > 1,000 parasites per microliter (estimated based on an average of 8000 leucocytes/ml) were treated in accordance with national guidelines. Asymptomatic *P. falciparum* gametocyte-positive children were recruited for the study. Gametocytic blood was collected by venipuncture in heparinized tubes. DMFA was performed with serum replacement as previously described [50]. Venous heparinized blood samples were centrifuged at 1300 *g* for 3 minutes. The natural plasma was then removed and the RBC pellet mixed with serum from a non-immune European AB blood donor. Mosquitoes held in 500 ml paper cups at a density of 80 per cup were allowed to feed on this blood for one hour. Non-fed or partially fed females were removed and discarded, while the remaining fully-engorged mosquitoes were kept in a biosafety room under the same standard conditions (12:12 LD, 27 ± 2°C, 70 ± 5% relative humidity).

### Experiment 1: Effects of age on mosquito competence

Three batches of about 1000 eggs each were collected from the *An. coluzzii* outbred colony at 4 days interval to generate three age classes of 4 days difference in the progeny (Figure 1). Resulting nymphs were collected over 2 successive days and placed in one of three emergence cages (30 x 30 x 30 cm) corresponding to each of the three age classes. Males were removed from the emergence cages after a period of 4 days, such that females in each age class were exposed to males for 3-4 days, thus ensuring an equivalent insemination rate among the three age classes. In order to respect natural mosquito gonotrophic cycles, mosquito females were blood-fed on rabbit every 4 days and oviposition cups were placed in the cages (Figure 1). From emergence to infection, mosquitoes also had *ad libitum* access to a 5% glucose solution. On the day of infection, mosquitoes were thus either (i) 3-4 day-old (received only one infectious blood meal), 7-8 day-old (received one blood meal on a rabbit and one infectious blood meal) and 11-12 days-old (received two blood meals on a rabbit and one infectious blood meal). Hereafter, these three age classes are named 4; 8 and 12 day-old respectively. Experimental infections were performed using a total of four parasite isolates over two distinct replicates (replicate 1: isolates A with gametocytemia of 40 gam µl^−1^ of blood and B, 32 gam µl^−1^; replicate 2: isolates C 80 gam µl^−1^ and D 40 gam µl^−1^). At day 8 post-infectious blood meal (dpibm), 50 mosquitoes from each age class and isolate (50 mosquitoes * 3 age groups * 4 parasite isolates) were dissected to count the number of developing oocysts in the midgut of infected females. At 14 dpibm, 50 mosquito head/thoraces from each age class and isolate (50 mosquitoes * 3 groups * 4 parasite isolates) were stored at −20 °C for further qRT-PCR (see below) to detect the presence of sporozoites. Four traits characterizing vector competence for *P. falciparum* were measured: (i) oocyst prevalence, expressed as the proportion of mosquitoes exposed to the infectious blood meal and harboring at least one oocyst in their midgut at 8 dpibm, (ii) oocyst intensity, expressed as the mean number of developing oocysts in the guts of infected females at 8 dpibm, (iii) sporozoites prevalence, expressed as the proportion of mosquitoes exposed to the infectious blood meal and having disseminated sporozoites in their head/thoraces at 14 dpibm, and (iv) sporozoites intensity, expressed as the estimated number of sporozoites. This number was computed by using the following transformation: 10^−((Ct-40)/3.3)^. This transformation considers the maximum number of amplification cycles (Ct) used in the qRT-PCR process, ensuring that the probability of detecting sporozoites becomes zero at higher amplification cycles (Ct=40). The coefficient 3.3 is derived from the negative slope observed in the linear relationship between the number of amplification cycles and the logarithm of the sporozoite density[51].

**Figure 1:**
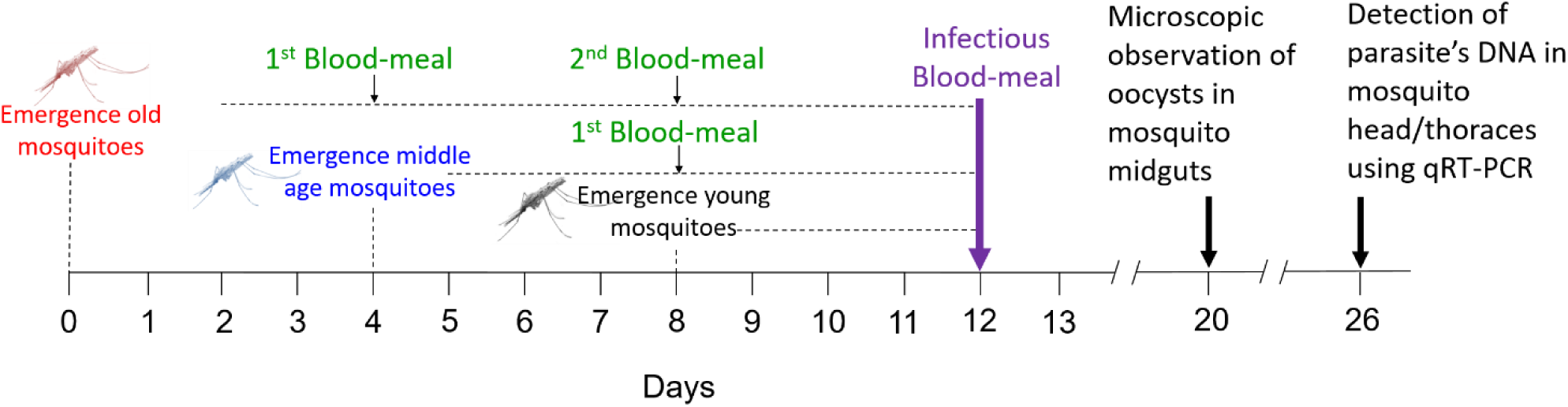
schematic representation of the design used in experiments 1 and 2. Mosquitoes belonging to one of three-age class (red: old, blue: middle age, and black: young mosquitoes) were challenged with one of 4 parasite isolates 12 days following the emergence of the old mosquito batch. There was a four days difference among the 3 age classes. When mosquitoes received the infectious blood-meal on day 12 (purple font), the old batch was 12 day-old, the middle age batch 8 day-old and the young batch 4 day-old. On day 20, corresponding to 8 days post infectious blood meal, mosquitoes were dissected to count the number of developing oocyst in their midgut. On day 26, corresponding to 14 days post infectious blood meal, the presence of parasite in mosquito head/thorax was assessed using qRT-PCR. On day 12 (infectious blood-meal), 12 day-old mosquitoes were not only older than 8 and 4 day-old mosquitoes, they also received more bloodmeals from rabbits (green font). The old batch received a total of three blood meals, middle age 2 bloodmeals and young a single infectious bloodmeal.

### Experiment 2: Effects of age and infection on mosquito survival

This experiment was conducted to explore possible age-mediated effect of infection on mosquito survival. Following the same procedure as described above*, An. coluzzii* mosquitoes belonging to one of three age classes (4, 8 or 12 day-old) were fed with either a gametocyte-positive (infected) or gametocyte-negative (control) blood. The same wild parasite isolates as in the first experiment (i.e. isolates A to D) were used. Uninfected control mosquitoes were provided with gametocytic blood that had been subjected to heat treatment to kill gametocytes. Half of the venous blood drawn from each carrier was heated at 45°C for 20 minutes. This heat treatment eliminates the gametocytes and has no impact on the survival and fecundity of female mosquitoes [52]. Since the nutritive quality of blood can significantly vary among individuals, particularly between infected and uninfected individuals, this procedure allows experiments to avoid potential confounding effects arising from different blood sources on the performance of infected and control mosquitoes. A total of 295 blood-fed control and 362 blood-fed parasite-exposed mosquitoes were obtained and were distributed in 500 ml paper cups (4 parasite isolates * 3 age classes * 2 infection status * 2 paper cups to avoid cup effect = 48 paper cups of ∼15 mosquitoes (range 4 to 18) each). Mosquito survival was recorded daily and dead mosquitoes from the infectious blood group were individually stored at −20°C for further PCR to assess their infection status [53]. Following PCR on the carcass of these dead individuals, three infection status were obtained: (i) uninfected control mosquitoes fed on heat-treated blood, (ii) exposed-infected mosquitoes fed on gametocyte-positive blood and (iii) exposed-uninfected mosquitoes fed on gametocyte-positive blood but in which the parasite failed to establish.

### Experiment 3: Effects of mosquito age on the parasite’s Extrinsic Incubation Period (EIP)

To assess the EIP, the 4-day-old and 12-day-old age classes only were specifically selected for this experiment, given the need for a substantial mosquito population and daily dissections to detect the presence of the parasite. Experimental infections of 4-day-old and 12-day-old mosquitoes were performed using a total of three parasite isolates over two distinct replicates (replicate 1: isolates E with 64 gametocytes/µl and F with 72 gametocytes/µl of blood; replicate 2: isolate G with 80 gametocytes/µl). Mosquitoes were kept at 27 °C and 75 % relative humidity, and the extrinsic incubation period of *P. falciparum* was examined through microscopic observation of mosquito gut and salivary glands from 8 to 16 dpibm. First, at 6 dpibm, between 30 and 32 mosquitoes of each age class and infected with isolate E or F were dissected to verify the success of infection and confirm the competence results obtained as part of experiment 1. Second, from 8 to 16 dpibm, 10 to 20 mosquito females (median=13) from each age class and each of the three isolates were dissected daily (N total=624). The presence and number of oocysts in mosquito guts and the presence of sporozoites in the salivary glands were assessed microscopically. Oocyst rupture in mosquito midgut and sporozoite invasion of salivary glands is highly asynchronous: while some oocysts are intact and keep developing within a given mosquito gut, others have already ruptured and released their sporozoites [54,55]. To estimate EIP, three metrics were derived from the microscopic observations:

(i) the proportion of infected mosquitoes with ruptured oocysts at 8-14 dpibm. This is the number of mosquitoes with at least one ruptured oocyst in their midguts out of the total number of infected mosquitoes (i.e. harboring either intact and/or ruptured oocysts);
(ii) the fraction of ruptured oocysts at 8-14 dpibm. This is, for each infected mosquito, the number of ruptured oocysts out of the total number of oocysts (intact + ruptured);
(iii) the proportion of oocyst-infected mosquitoes with sporozoites in their salivary glands at 8-14 dpibm. This is the number of oocyst-infected mosquitoes harboring sporozoites in their salivary glands out of the total number of infected mosquitoes (i.e. harboring either intact and/or ruptured oocysts).

### qPCR detection of Plasmodium falciparum

The detection and quantification of parasite’s DNA in mosquitoes dissected at 14 dpibm (experiment 1) was performed by real-time PCR using standard procedures [56]. The mitochondrial gene that codes for the cytochrome C Oxidase (Cox1) was targeted. The sequences of the primers used were: qPCR-PfF 5’-TTACATCAGGAATGTTATTGC-3 ‘and qPCR-PfR 5’-ATATTGGATCTCCTGCAAAT-3’. Sample reaction occurred in a total volume of 10 μl containing 1 μl of DNA (∼ 40 ng/μl); 4.6μL of water; 2μl of 1x HOT Pol Eva Green qPCR Mix Plus ROX and 1.2μl of each primer at 5μM. The amplification began with an activation step at a temperature of 95 °C for 15 min and 40 cycles of denaturation at 95 °C to 15 s, annealing / extension at 58 °C for 30 s.

### Statistics

Experiment 1. The effect of mosquito age class (three levels: 4-, 8- or 12-day-old) on oocyst and sporozoite prevalence was examined using a binomial Generalized Linear Mixed Model (GLMM, glmmTMB package [57]). Oocyst and sporozoite intensity were analysed using a zero-truncated negative binomial GLMM and a gaussian GLMM, respectively (glmmTMB package). In these GLMMs, age class was set as a fixed effect and parasite isolates (A to D) as a random effect. Experiment 2. The effect of mosquito age class and infection status (three levels: uninfected control, exposed-infected and exposed-uninfected) on mosquito survival was explored using a mixed effect Cox’s proportional hazard regression models (coxme package [58]). Mosquito age, infection status and their interaction were considered as fixed effects whereas cup identity and parasite isolate were considered as random effects. Experiment 3. Binomial GLMMs were used to test the effect of age class (two levels: 4 or 12 day-old), dpibm and the interaction on (i) the proportion of infected mosquitoes with ruptured oocysts, (ii) the fraction of ruptured oocysts, and (iii) the proportion of oocyst-infected mosquitoes with sporozoites in their salivary glands. In these GLMMs, age class and dpibm were set as a fixed effects and parasite isolates (F to H) as a random effect. the “Anova” function from the “car” library version 3.1-1 [59] was used to estimate the significance of terms. When age was significant, multiple pairwise post-hoc tests were performed to compare age classes using the “emmeans” library [60].

### Modeling the consequence of mosquito age variation on malaria transmission

Based on the age-structured model for the human-mosquito transmission proposed and analyzed in [46], the vectorial capacity (*V*_*C*_ (*a*)) of a mosquito aged *a* is given by :

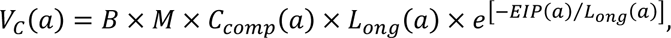

where *B* is the mosquito biting rate, *M* is the mosquito to human ratio, and *L*_*ong*_(*a*), *EIP*(*a*) and *C*_*comp*_(*a*) are longevity, extrinsic incubation period, and competence of a mosquito of age *a* respectively. Mosquito competence (*C*_*comp*_(*a*)) encompasses other quantitative traits expressed during the basic steps of the parasite infection events which are, the infection probability (aka prevalence) (*P*_*inf*_(*a*)), the number of developing oocysts (*N*_*oocysts*_(*a*)) (aka intensity), and the conversion probability from oocysts to sporozoites (*P*_*oocyst*→*spo*_(*a*)) of a mosquito of age (*a*). Consequently, we have *C*_*comp*_(*a*) = *P*_*inf*_(*a*) × *N*_*oocysts*_(*a*) × *P*_*oocyst*→*spo*_(*a*). We simulated the expected vectorial capacity *V*_*C*_ (*a*) for mosquitoes belonging to age class 4-day-old vs. 12-day-old. We explored the parameter space through a Latin hypercube sampling with 10,000 replicates for which all parameters were randomly chosen within their confidence interval based on the data measurements obtained experimentally here. See parameter values in Table S1.

## Results

### Experiment 1: Effects of age on mosquito competence

Twelve-day-old mosquitoes showed a lower likelihood of harboring oocysts at 8 dpibm compared to both 8-day-old and 4-day-old mosquitoes (46.7 ± 7% vs 60 ± 7% and 61 ± 7% in 8 day-old and 4 day old, respectively; LRT *X^2^*_2_ = 10.6, P=0.005, Figure 1A). Additionally, the number of oocysts in the mosquito midgut (± se) was significantly influenced by mosquito age (LRT *X^2^*_2_ = 8.8, P=0.01), with 4-day-old mosquitoes having a higher oocyst count compared to 12-day-old mosquitoes (4, 8 and 12 day-old mosquitoes carried on average 4.57 ± 0.32 oocysts, 3.66 ± 0.26 and 3.3 ± 0.38, respectively, Figure 1B). A similar trend was observed at 14 dpibm, where 12-day-old mosquitoes exhibited both reduced sporozoite prevalence (54.5 ± 7% vs 74 ± 5% and 73 ± 5%, LRT *X^2^*_2_ = 30, P<0.0001, Figure 1C) and intensity (LRT *X^2^*_2_ = 49.8, P<0.0001, Figure 1D). Detailed results on infection prevalence and intensity for each parasite isolate (A to D) are given in Figure S1.

### Experiment 2: Effects of age and infection on mosquito survival

The survival of infected and uninfected mosquitoes belonging to the three age classes was monitored daily. Overall, mosquito age significantly influenced mosquito survivorship (LRT *X^2^*_2_ = 48, P < 0.0001, Figure 3). Twelve day-old mosquitoes had a median longevity of 18 days and died at a faster rate than both 8 (median longevity of 21 days) and 4 day-old mosquitoes (median longevity 23 days). All three pairwise post-hoc comparisons were significant (12 vs 8 day-old: z = 4, P < 0.001; 12 vs 4 day-old: z = 7, P< 0.001, 8 vs 4-day-old: z = 3, P = 0.005). There was no significant main effect of mosquito infection status on overall survival (LRT *X^2^*_2_ = 0.39, P = 0.82). However, the impact of infection on survival varied across the three age classes, as indicated by a significant interaction between infection status and age class (LRT *X^2^*_4_ = 15.8, P = 0.003). Specifically, while no effect of infection status on the survival of 4- or 8-day-old mosquitoes was observed, 12-day-old infected mosquitoes had a significantly longer lifespan compared to their uninfected counterparts of the same age (z = 2.7, P = 0.007, Figure 3; Figure S2). *P. falciparum* infection in 12-day-old mosquitoes extended the median survival by 2 days compared to uninfected controls and by 8 days in exposed-uninfected individuals.

**Figure 2:**
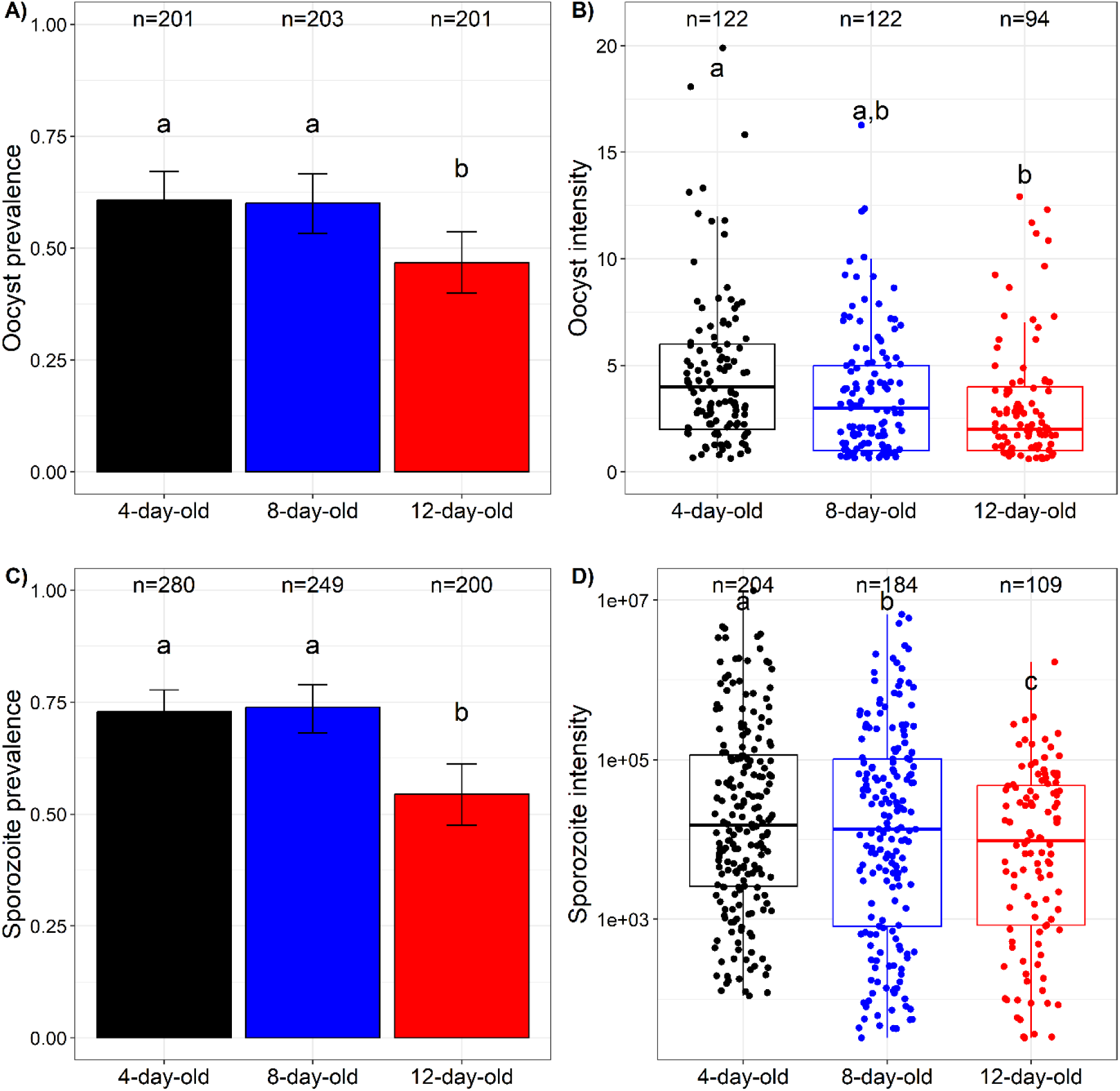
Effects of mosquito age on competence for *Plasmodium falciparum.* (A) Oocyst prevalence, expressed as the proportion of mosquitoes exposed to an infectious blood meal and harboring at least one oocyst in their midgut at 8 dpibm. (B) Oocyst intensity, expressed as the number of developing oocysts in the guts of infected females at 8 dpibm. (C) Sporozoites prevalence, expressed as the proportion of mosquitoes exposed to an infectious blood meal harboring disseminated sporozoites in their head/thoraces at 14 dpibm. (D) Sporozoites intensity, expressed as the estimated number of sporozoites. The y-axis in this panel is on a log10 scale. Two experimental replicates were performed each with two distinct wild parasite isolates. n = number of dissected mosquitoes (about 50 for each of the four parasite isolate). Different letters denote statistically significant differences based on multiple pair-wise post-hoc tests.

**Figure 3:**
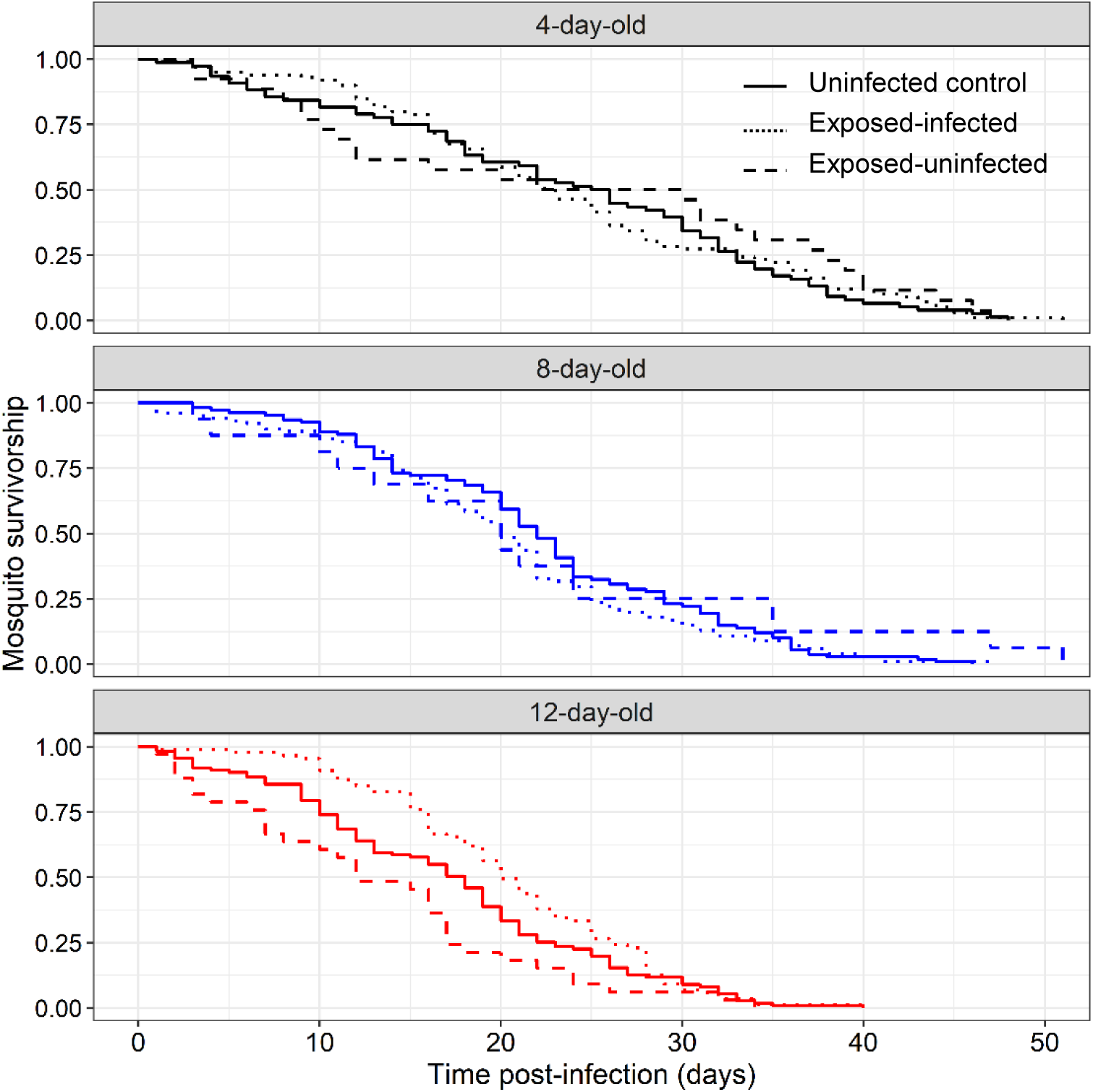
Effects of mosquito age and infection status on survival. Survival was recorded daily from day 1 post infectious bloodmeal until the death of all mosquitoes from a total of 48 paper cups containing circa 15 mosquitoes each. The sample size of uninfected control, exposed-infected and exposed-uninfected mosquitoes were 76, 99, 26 (4-day-old); 108, 101, 16 (8-day-old); and 111, 87, 33 (12-day-old) respectively. N total = 657. Solid, dashed and dotted lines show uninfected control, exposed-uninfected and exposed-infected mosquitoes, respectively.

### Experiment 3: Effects of age on the parasite’s Extrinsic Incubation Period (EIP)

The development of *P. falciparum* was monitored in two age classes (4 and 12 day-old at the time of infection) over a period of 8 to 16 dpibm using three different parasite isolates (E, F, and G). Microscopic examination of oocysts at 6 dpibm confirmed that old mosquitoes exhibited lower competence for *P. falciparum* compared to their young counterparts, as indicated by both lower oocyst prevalence and intensity (Figure S3). By 15 dpibm, all mosquitoes exposed to the G isolate had died. Therefore, the subsequent analysis focused on the period between 8 to 14 dpibm for all three isolates. The results for each isolate separately until 17 dpibm are detailed in Figure S4. Uninfected mosquitoes (n=39 individuals from isolate H, n=41 from isolate I, and n=106 from isolate J), from which no estimates of EIP could be derived, were excluded from the analysis.

None of the infected mosquitoes exhibited ruptured oocysts at 8 dpibm and the first observations occurred at 9 dpibm for both age class. As expected, there was a highly significant positive relationship between time post-bloodmeal and the proportion of mosquitoes showing ruptured oocysts (*LRT X^2^_1_* = 95, P < 0.0001, Figure 4A). The timing of rupturing was similar between the two age classes (no dpibm by mosquito age interaction: *LRT X^2^_1_* = 0.2, P = 0.66, Figure 4A). Using this metric, the estimated EIP_50_ from the binomial models were 11.90 days for 12-day-old mosquitoes and 12 days for 4-day-old mosquitoes.

**Figure 4:**
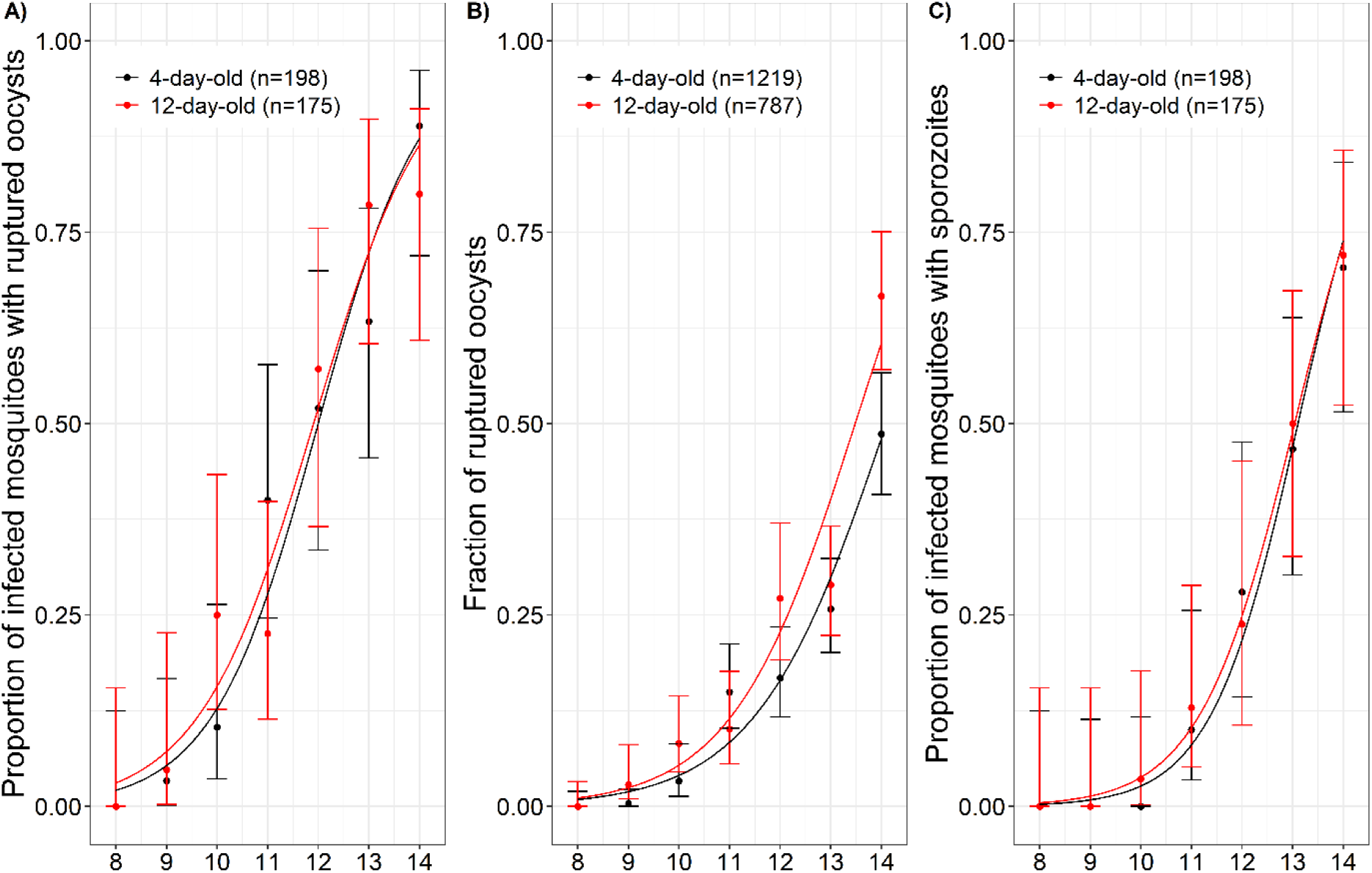
Effects of mosquito age on the parasite extrinsic incubation period. A) Proportion of infected mosquitoes with ruptured oocysts (± 95% CI) from 8 to 14 dpibm, expressed as the number of mosquitoes with at least one ruptured oocyst out of the total number of infected mosquitoes (i.e. harboring either intact and/or ruptured oocysts) for two age classes (4-day-old in black and 12-day-old mosquitoes in red). B) Fraction of ruptured oocysts (± 95% CI), expressed as the number of ruptured oocysts out of the total number of oocysts (intact + ruptured). C) Proportion of infected mosquitoes with sporozoites in their salivary glands (± 95% CI), expressed as the number of oocyst-infected mosquitoes harboring sporozoites in their salivary glands out of the total number of infected mosquitoes. The lines represent best-fit logistic regression curves for each age class. Three parasite isolates (E, F, G) were used over two experimental replicates. A) to C): Sample size = 21 to 31 mosquito /day/age class (median=27.5). In panels A) and C) the sample sizes indicate total mosquito numbers while in in panel C) total oocyst number.

The fraction of ruptured oocysts in infected mosquitoes tended to be higher in 12-day-old compared to 4-day-old mosquitoes, although this was not statistically significant (12-day-olds: 160 ruptured oocysts out of 787 oocysts counted (20%) between 8 and 16 dpibm, 4-day-olds: 177 / 1 219 (14.5%) over the same period, LRT *X^2^_1_* = 1.3, P = 0.26, Figure 4B). This trend seemed to be driven by the density of oocysts rather than the age of the mosquitoes: older mosquitoes exhibited lower oocyst densities compared to younger mosquitoes (Figure S5A), and a negative association was observed between oocyst density and the proportion of ruptured oocysts (Figure S5B).

Finally, the proportion of infected mosquitoes with disseminated sporozoites in their salivary glands, the most epidemiologically-relevant metric, did not vary between the two age classes (LRT *X^2^_1_* = 0.03, P = 0.87, Figure 4C). The proportion of sporozoite-infested salivary glands increased with dpibm similarly for both age class (i.e. no significant dpibm by age interaction, LRT *X^2^_1_* = 0.2, P = 0.65). Using this metric, the estimated EIP_50_ from the binomial models were 13.04 days for 12-day-old mosquitoes and 13.10 for 4-day-old mosquitoes.

The estimated time for sporozoites to migrate and invade mosquito salivary glands following the egress from the oocysts was 27.36 hours for 12-day-old mosquitoes (*i.e.*, the difference between the observed sporozoite presence in salivary glands (Fig. 4C) and the time it takes for oocyst rupturing (Fig. 4A): 13.04 – 11.90 days). For 4-day-old mosquitoes, this time was 26.4 hours (13.10 –12 days).

### Modeling the consequence of mosquito age variation on malaria transmission

To investigate the epidemiological impact of mosquito age on vectorial capacity, we built a mathematical model based on the experimental values of mosquito competence, survival and the parasite EIP obtained in this study for mosquitoes belonging to age class 4-day-old vs 12-day-old. Mosquito competence encompassed the oocyst prevalence, intensity and the conversion probability from oocysts to sporozoites (*P*_*oocyst*→*spo*_(*a*)) of a mosquito of age (*a*). This parameter (*P*_*oocyst*→*spo*_) turned out to be a nonlinear decreasing function with respect to the number of developing oocysts (*N*_*oocysts*_) (Figure S5), and we therefore considered 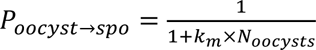, where *k*_*m*_ is the fitted parameter.

Overall, the mean (± se) vectorial capacity of the young 4-day-old mosquitoes was 107.75 ± 0.84, significantly exceeding that of the 12-day-old counterparts, which had a *V*_*C*_ of 76.69 ± 0.42 (Figure 5, scenario 1, LRT *X^2^_1_* = 1203, P <0.001). To assess the epidemiological consequences of the positive effect of infection on the survival of 12-day-old mosquitoes (figure 3), we explored two other scenarios for which the survival values observed in infected 12-day-old was replaced by that of either 12-day-old controls (scenario 2), or exposed-uninfected 12-day-old (scenario 3). The average *V*_*C*_ of 12-day-old mosquitoes stood at 59.90 ± 0.38 for scenario 2 and 41.79 ± 0.09 for scenario 3. Therefore, the positive impact of infection on the survival of 12-day-old mosquitoes amplified their *V*_*C*_ by 28% and 84% in comparison to scenarios 2 and 3, respectively.

**Figure 5:**
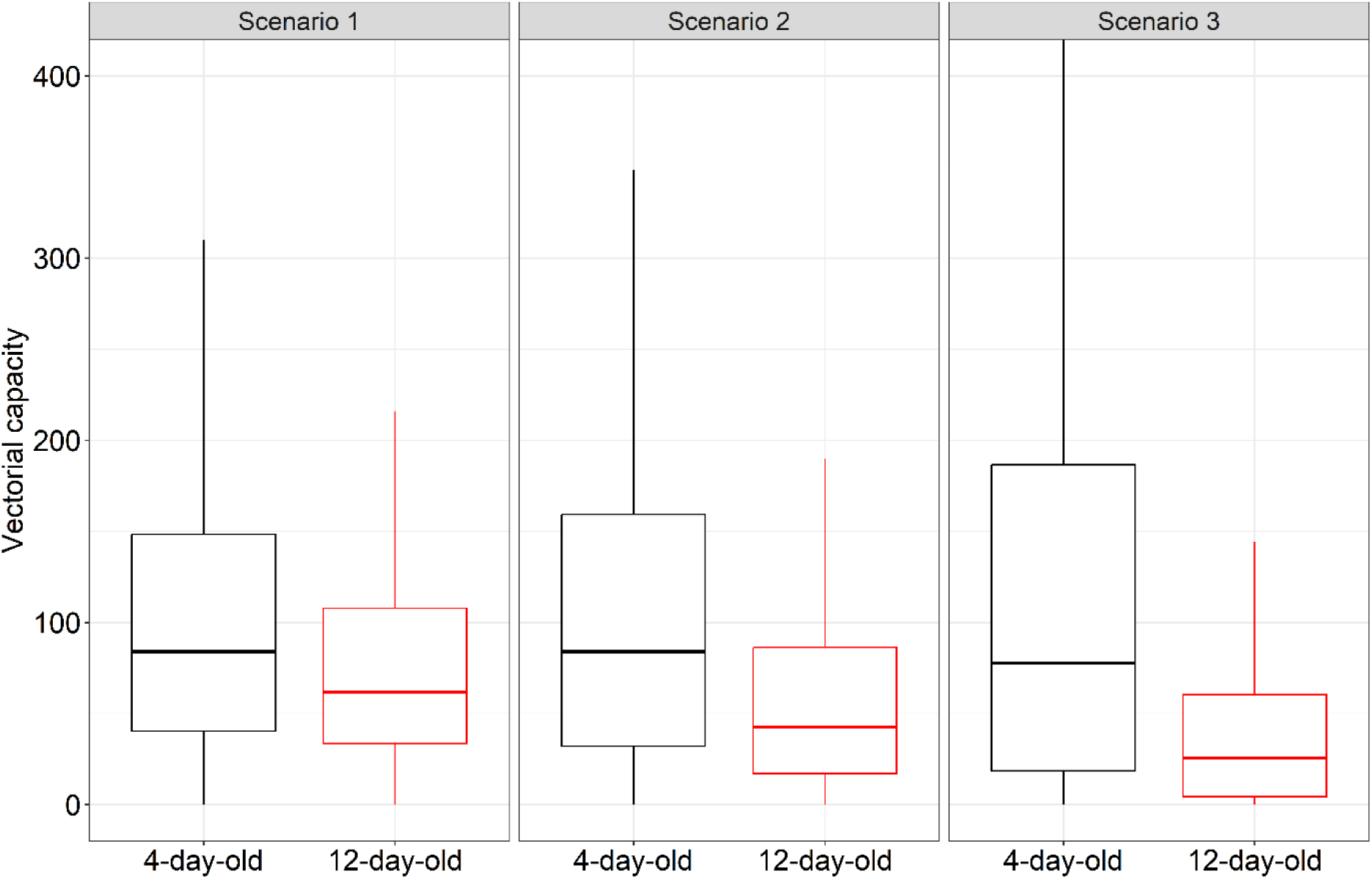
Effects of mosquito age on vectorial capacity. Scenario 1: predicted age-dependent model of vectorial capacity based on the survival, competence, and EIP data from this study. Scenario 2: same as scenario 1 except that the survival values of infected 12-day-old mosquitoes were replaced by that of 12-day-old uninfected controls. Scenario 3: same as scenario 1 except that the survival values of infected 12-day-old mosquitoes were replaced by that of 12-day-old exposed-uninfected individuals.

## Discussion

The findings presented in this study reveal age-related variations in key parameters influencing the transmission potential of *P. falciparum*, the causative agent of the most severe form of human malaria. Firstly, 12-day-old females displayed both increased qualitative and quantitative resistance, characterized by reduced parasite prevalence and intensity, respectively compared to younger females. Older mosquitoes therefore appear to be less competent hosts for this malaria parasite. Secondly, despite the accelerated mortality rate observed in older mosquitoes indicative of overall senescence, there was an age-mediated effect of infection with improved survival specifically among the infected 12-day-old mosquitoes. Thirdly, regardless of mosquito age, the median EIP remained consistent with a 13-day duration. Integrating these findings into an epidemiological model revealed that while the vectorial capacity of older mosquitoes was lower than that of younger ones, it remains substantial due to the extended longevity of old individuals in the presence of infection. Examining host-parasite interactions typically involves experimental infections of hosts of the same age, often young individuals, to monitor parasite development while minimizing loss of individuals over time. While this approach helps to minimize the confounding effects of host age, our findings strongly emphasize that the outcomes achieved with this practice cannot be extrapolated to the entire population.

The results of the first experiment contradict the immunosenescence hypothesis, which posits that the ability of insects to clear infections should diminish as they age [14–16]. Rather, the higher potential for parasite establishment and development observed in younger mosquitoes compared to the 12-day-old cohort highlights a declining vector competence with increasing mosquito age. These findings are consistent with previous studies that have explored the influence of mosquito age on their competence for *Plasmodium* parasites. Terzian et al. [61] similarly observed an age-related increase in resistance to *P. gallinaceum* in *Ae. aegypti*. Okech et al. [62] found no discernible effect of age on the prevalence of *P. falciparum* infections in *An. gambiae*; however, the extremely low mosquito infection rates (circa 5%) and intensity (circa one oocyst) across all age groups may have limited their statistical power. More recently, Pigeault et al.[39] provided evidence that, despite a decline in haemocyte numbers as the mosquito *Culex pipiens* ages, older individuals displayed significantly greater resistance to *P. relictum* infection compared to their younger counterparts.

The association between host age and competence is more ambivalent when considering insect vectors of pathogens other than *Plasmodium*. While tsetse flies, *Ae. aegypti*, and *Cu. tritaeniorhynchus* infected with a trypanosome, a filarial nematode, and a West Nile virus, respectively, demonstrated reduced competence in older cohorts [63–65], an inverse effect was observed in *Ae. albopictus* and *Ae. aegypti* infected with Zika and Sindbis virus, respectively [66,67]. Moreover, no age effect on infection with an entomopathogenic fungus was observed in *An. gambiae* [68], and the impact of *Cu. quinquefasciatus* age on West Nile virus infection varied depending on virus dose, mosquito strain, and temperature [69]. The proximate mechanisms underpinning age-related variations in competence remain unresolved and can vary among the different biological systems, but changes in molecular and physiological environment of the midgut such as microbiota composition could be involved [63].

Studies have shown that a non-infectious blood meal preceding the infectious blood meal offset the effect of age on competence [39,61,65]. The authors proposed that older females may experience depleted energy reserves after an extended period without blood feeding, and an additional blood meal could replenish the necessary nutrients for parasite development. In contrast, other studies indicated that prior blood-feeding can not only stimulate the mosquito immune system [40–42] but also accelerate the digestion of the infectious blood meal, leaving less time for the parasite to undergo zygotes to ookinetes to oocysts transitions [43]. In our study, 12-, 8-, and 4-day-old females at the time of infection received 2, 1, or no prior bloodmeals respectively. Blood-feeding induces the proliferation of bacterial populations within the mosquito midgut, which can directly and/or indirectly (through immune system stimulation), mitigate *Plasmodium* infection [70]. Thus, we postulate that the increased resistance of older mosquitoes to infection can be attributed to the presence of a more mature immune system, abundant microbiota and/or an accelerated digestion of the infectious blood meal due to the previous feeding episodes. Other mechanisms may also be involved and could deserve considerations as part of future experiments. For example, the possibility that the size of the infectious bloodmeals taken by older mosquitoes were smaller than that of younger females, thereby reducing the number of ingested gametocytes cannot be excluded (although an inverse pattern was in fact observed in tsetse flies and *Ae. aegypti*, where older individuals ingested on average larger meals [63,71]). Furthermore, it is possible that the longer pre-infection period in older females could have selected for the most robust individuals.

Mosquito age significantly influenced mosquito survivorship, with older mosquitoes having shorter median longevities compared to younger age classes. This observation aligns with previous studies highlighting the existence of senescence in mosquitoes [25,72]. In addition, upon further analysis of these survival data in a separate article developing an age-structured epidemiological model, it was revealed that the mortality rate within each age class displayed non-constancy, increasing with age in a manner that resembles a Gompertz function [23]. This provides additional evidence supporting the notion that mosquitoes do undergo senescence.

Importantly, the influence of infection on survival varied across the age classes. While no effect of infection on the survival of 4- or 8-day-old mosquitoes was observed, infected 12-day-old mosquitoes exhibited a significantly longer lifespan compared to their uninfected counterparts of the same age. This suggests that the presence of the parasite may modulate the aging process and alter survival dynamics in older infected mosquitoes specifically. The impact of malaria infection on mosquito survival remains equivocal and appears to be contingent upon various factors, including the natural vs. artificial nature of the host-parasite combination [73], mosquito species [74] and strains [75,76], parasite lineages [52,74,77] and environmental conditions [52,78–81]. Our study contributes an additional factor to this list by demonstrating that the impact of infection on mosquito survival can also be age-dependent. To our knowledge, this is the first evidence of a positive effect of infection by the human malaria parasite *P. falciparum* on the survival of its vector mosquitoes. The fact that this effect is age-dependent and only observed in older mosquitoes probably explains why this was not evidenced in earlier studies. Our results therefore shed light on a fountain of youth effect of infection for old mosquitoes that partially restores their vectorial capacity and may have significant effect on malaria transmission.

Presently, the mechanistic explanations for the extended lifespan observed in infected old individuals are lacking. Previous research has revealed that parasites can extend the lifespan of their hosts, including mosquitoes, by disrupting the trade-off between fecundity and longevity [76,82–84]. For example, Vézilier et al. [76] reported an increased mosquito longevity concomitant to a decreased fecundity in individuals infected with the avian malaria *P. relictum*. The simultaneous examination of both survival and fecundity in infected mosquitoes belonging to various age classes will be required to elucidate this possibility. In addition to the fecundity-longevity trade-off, research indicates that *Plasmodium* infection in mosquitoes may induce increased food intake, leading to elevated energy input [85–87]. This heightened energy acquisition may not only facilitates parasite development but also contributes to better mosquito conditions, potentially resulting in an extended lifespan. This could be particularly true given that there appears to be a non-competitive relationship regarding resource utilization between the mosquito and its parasite [88,89]. Consequently, we anticipate that infected older mosquitoes would exhibit a greater propensity for food uptake compared to their younger infected counterparts. In a recent study, we found that *An. gambiae* exposed to wild isolates of *P. falciparum* in Burkina Faso but which remained uninfected exhibited reduced mosquito survival compared to successfully infected mosquitoes, although a non-exposed mosquito control group was not included in this study [90]. Here, we provide confirmation that resisting parasite infection can impose a survival cost on mosquitoes and that the extent of this cost can be age-dependent.

The sporogony represents the longest replicative process in the life cycle of malaria parasites. Given the considerable duration of this period relative to the average lifespan of the vector, we hypothesized that *P. falciparum* might possess the ability to plastically accelerate its developmental rate when its transmission potential is jeopardized by the advanced age of its mosquito vector. Similar condition-dependent developmental strategies have been documented in various parasite species [91], including blood-stage malaria parasites [92]. Contrary to expectations, our results revealed no significant variation in the timing of sporozoite appearance between 12 day-old and 4 day-old mosquitoes. Our observations align with a previous study that has also reported a lack of age-dependent difference in the EIP of the Zika virus in *Ae. aegypti* [93]. This emphasizes the consistency of EIP in the context of mosquito age and further suggest that the beneficial effects of lengthy EIPs must be under strong selective pressures [94,95]. That being said, recent studies have demonstrated that providing one or more non-infectious blood meals after the initial infectious meal significantly shorten the EIP [29,31,96–98]. It is common practice in laboratory studies on mosquito-*Plasmodium* interactions to maintain specimens on a sugar solution alone to prevent the decrease in sample size associated with each blood-feeding episode (non-fed individuals being excluded). Therefore, it would be interesting to test whether the EIP could vary with mosquito age when their gonotrophic cycles are respected following infection. Since the EIP is highly temperature-sensitive, it would also be intriguing to investigate whether the parasite could influence the thermal preference of older infected mosquitoes in a way that shorten the EIP.

In the context of the ongoing global malaria elimination efforts, there is a need to enhance our understanding of the parasite and vector components that influence transmission intensity. The age at which a mosquito acquires the parasite is expected to have a significant impact on its contribution to transmission. Older mosquitoes are less likely to transmit pathogens because they are less likely to survive during and after the EIP. Moreover, we found that the vectorial competence of old mosquitoes is reduced, providing additional evidence for their poor vectorial capacity. However, the observed parasite-induced extended lifespan in older mosquitoes tends to complicate this notion. Our modeling approach indicates that the capacity of older mosquitoes, while admittedly lower than that of younger mosquitoes, remains significant. The laboratory setting in which our experiments were conducted likely provided optimal survival conditions, raising concerns about the ecological relevance of the findings. It is crucial to investigate whether the observed extended survival of older infected mosquitoes holds true in natural field conditions, where various hazards may limit their lifespan. Furthermore, previous studies have demonstrated that mosquitoes carrying sporozoites may exhibit an elevated biting rate, which can contribute to higher feeding-mortality in natural environments.

The results of this study collectively emphasize the importance of considering mosquito age variation when examining disease transmission dynamics and open up a new perspective on the evolution of within-vector virulence. Ageing leads to physiological differences among mosquitoes, which in turn can exert selective pressures on their parasites. This selection process may eventually result in parasite adaptations that are specific to the most commonly encountered host age or in the differential expression of parasite traits depending on the age of the host in which they reside. To transfer our findings to natural settings, it will be crucial to enhance the accuracy of existing tools for classifying the age of wild mosquitoes [99].

