## supplementary file for "Mosquito aging modulates the development, virulence and transmission potential of pathogens"

**Supplementary information**

Table S1: Parameters used in the mathematical model.

| Parameter, (description), (unit) | Mosquito age in days | |
|---|---|---|
| | 4-day-old | 12-day-old |
| $k_m$ , (fitted parameter of the conversion probability from oocysts to sporozoites), (/oocyts) | 0.212 | 0.160 |
| $P_{inf}(a)$, (infection probability), (unitless) | 0.61 | 0.47 |
| EIP, (extrinsic incubation period), (days) | 13.10 | 13.04 |
| $B$, (biting rate), (days.ind$^{-1}$) | 1/3 | 1/3 |
| M, (mosquito to human ratio), (unitless) | 20 | 20 |

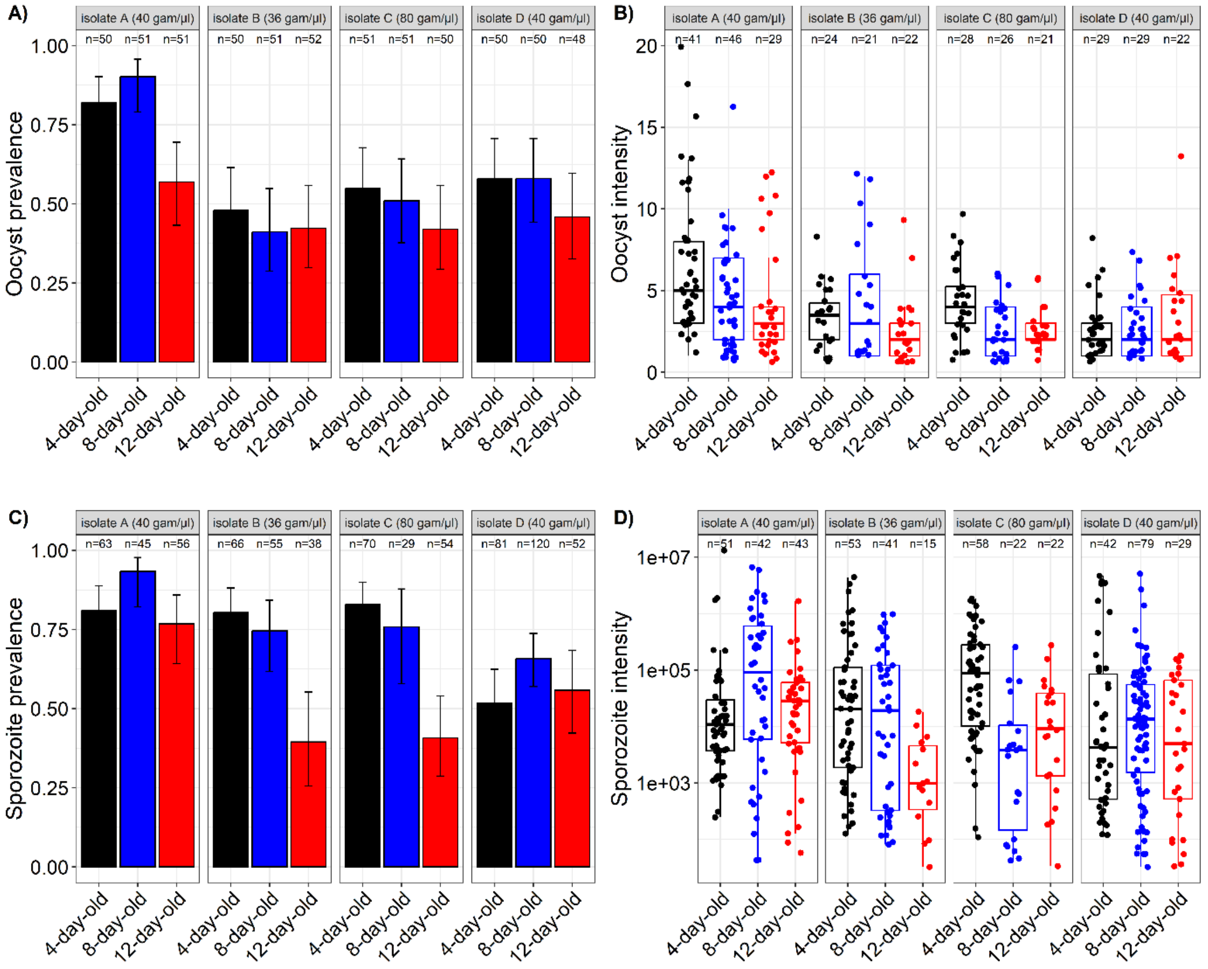


**Figure S1 : Oocyst prevalence and intensity in *Anopheles coluzzii* belonging to three age classes (4 day-old vs 8 day-old vs 12 day-old on the day of infection)**. (A) Oocyst prevalence is expressed as the proportion of mosquitoes exposed to an infectious blood meal and harboring at least one oocyst in their midgut at 8 dpi. (B) Oocyst intensity is expressed as the mean number of developing oocysts in the guts of infected females at 8 dpi. (C) Sporozoite prevalence is expressed as the proportion of mosquitoes exposed to an infectious blood meal and harboring disseminated sporozoites in their head/thoraces at 14 dpi. (D) Sporozoite intensity, expressed as the estimated number of sporozoites. Data are shown for each parasite isolate: A, B, C and D. n = number of dissected mosquitoes.

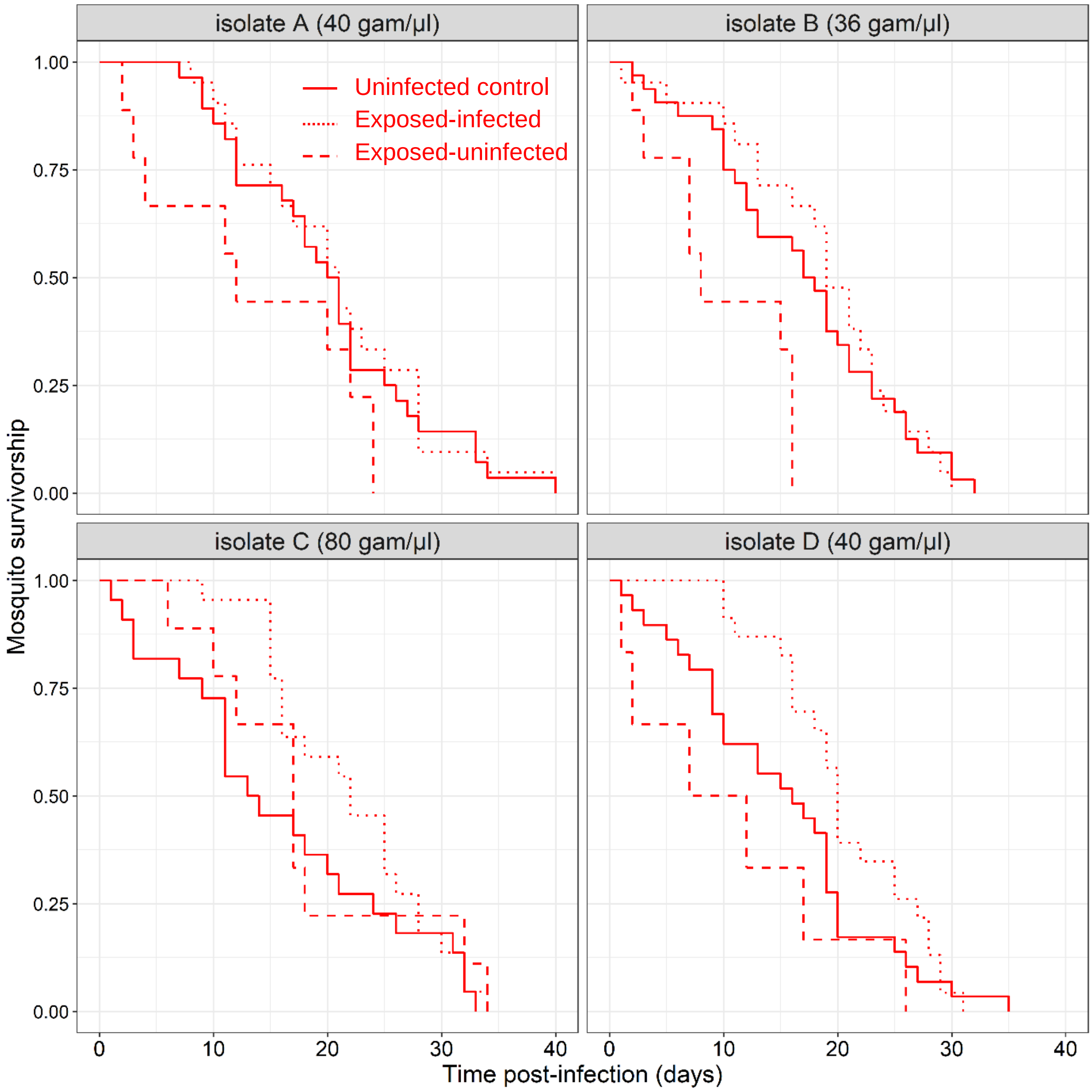


Figure S2: **Survivorship of 12 day-old mosquitoes as a function of infection status for each of the 4 parasite isolates (A, B, C, D).** The sample size of uninfected control, exposed-infected and exposed-uninfected were 28, 21, 9 (isolate A); 32, 21, 9 (isolate B); 22, 22, 9 (isolate C) and 29, 6, 23 (isolate D).

**FIGURE S3**

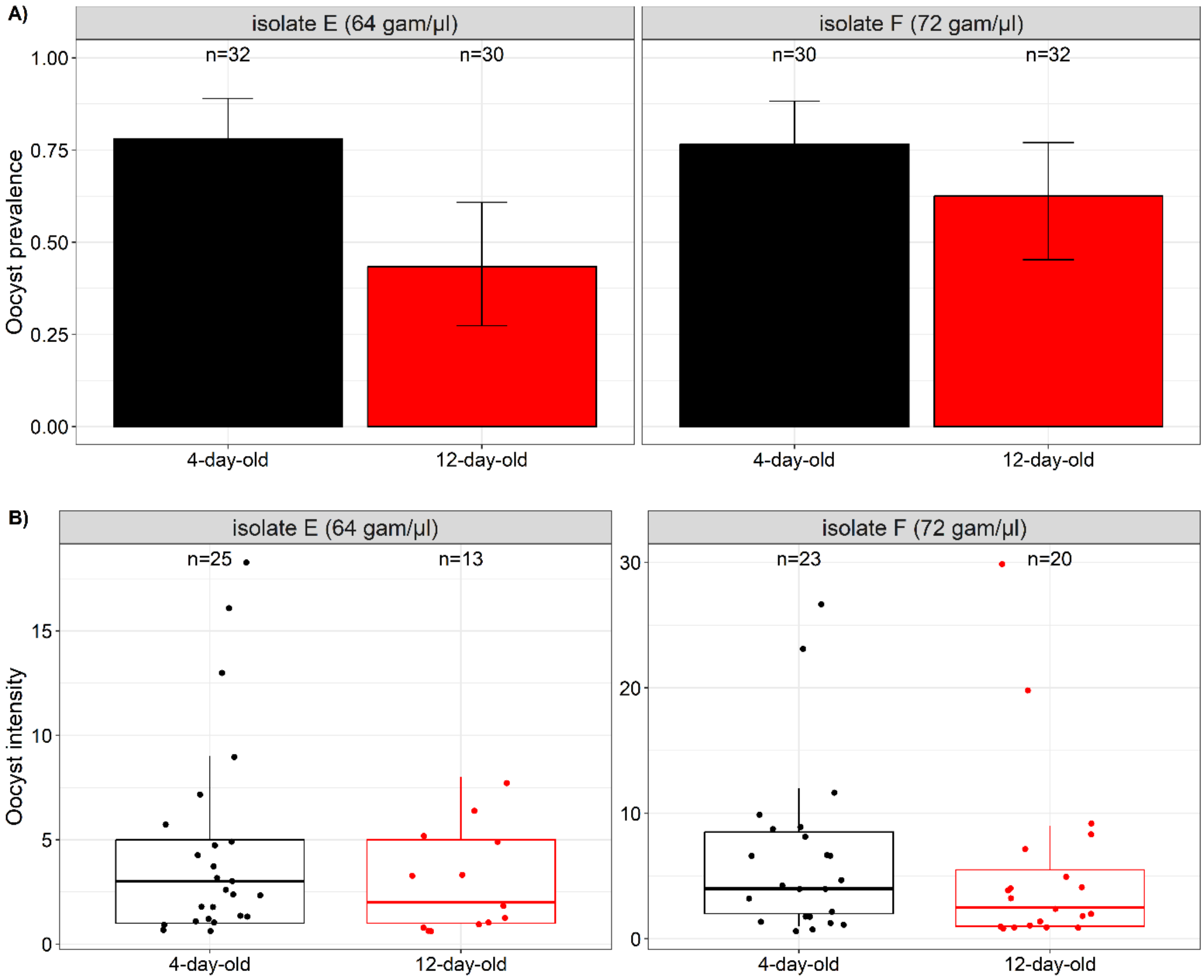


**Figure S3: Effects of age on mosquito competence for *Plasmodium falciparum*:** Oocyst prevalence (± 95% CI) on day 6-9 post blood meal (dpbm), expressed as the number of mosquito females harboring at least one oocyst in their midguts out of the total number of dissected females, and oocyst density at 6-9 dpi, expressed as the mean number of developing oocysts in the guts of infected females. N = number of dissected mosquitoes over 2 experimental replicates and using a total of 3 parasite isolates. Different letters denote statistically significant differences based on multiple pair-wise post-hoc tests.

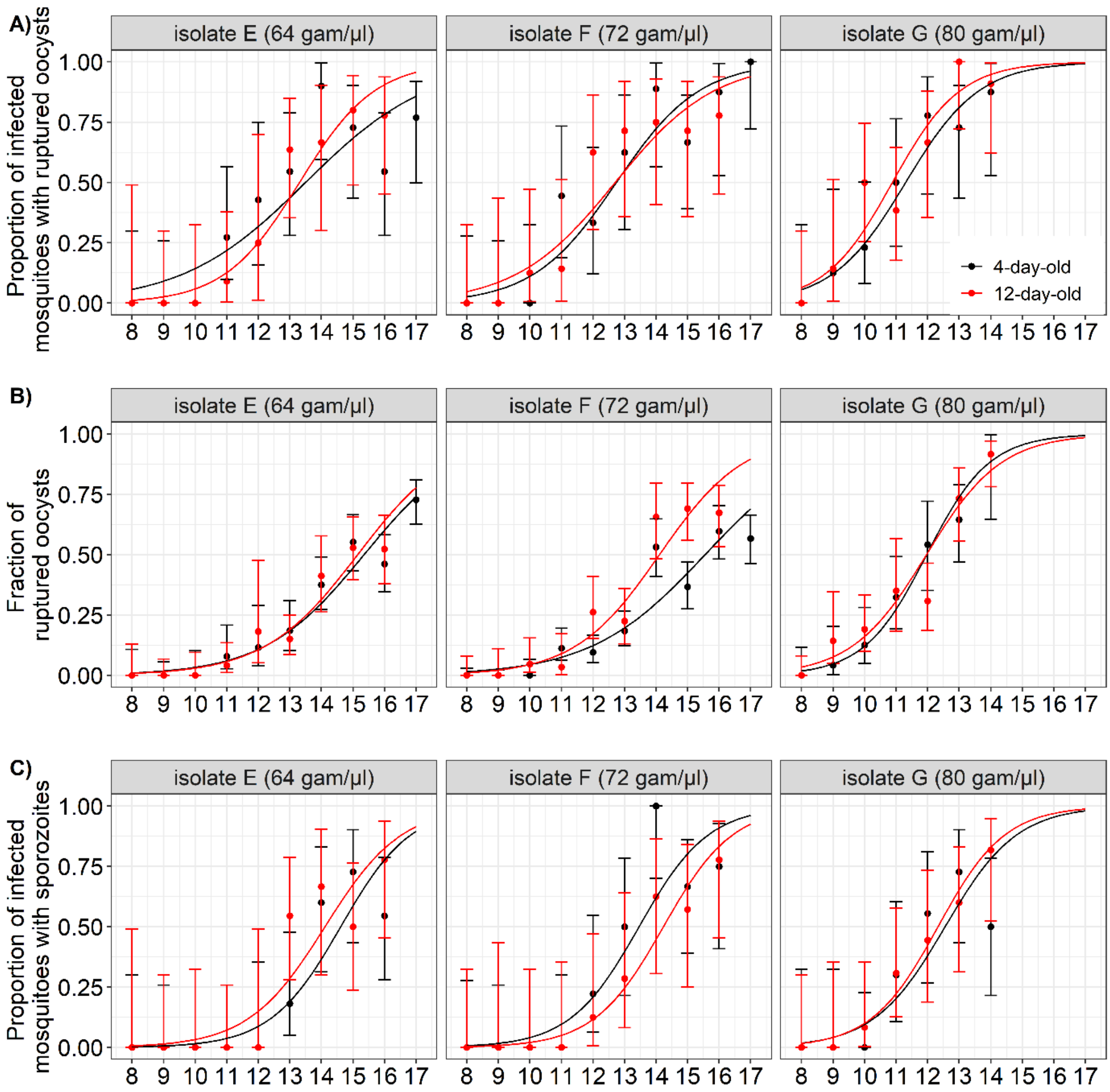


**Figure S4: Effects of mosquito age on the parasite's Extrinsic Incubation Period (EIP)**. A) Proportion of infected mosquitoes with ruptured oocysts (± 95% CI) from 8 to 17 dpbm, expressed as the number of mosquitoes with at least one ruptured oocyst out of the total number of infected mosquitoes (i.e. harboring either intact and/or ruptured oocysts) for two age classes (4-day-old in black and 12-day-old mosquitoes in red) and 3 parasite isolates (E, F, G). B) Fraction of ruptured oocysts (± 95% CI), expressed as the number of ruptured oocysts out of the total number of oocysts (intact + ruptured) for the two age classes and the 3 parasite isolates (C) Proportion of oocyst-infected mosquitoes with microscope-identified sporozoites in the salivary glands (± 95% CI), expressed as the number of oocyst-infected mosquitoes harboring sporozoites in their salivary glands out of the total number of infected mosquitoes, for the two age classes and the 3 parasite isolates. The lines represent best-fit logistic regression curves for each isolate. A to C: Sample size = 10 to 12 midguts/day/isolate/age class.

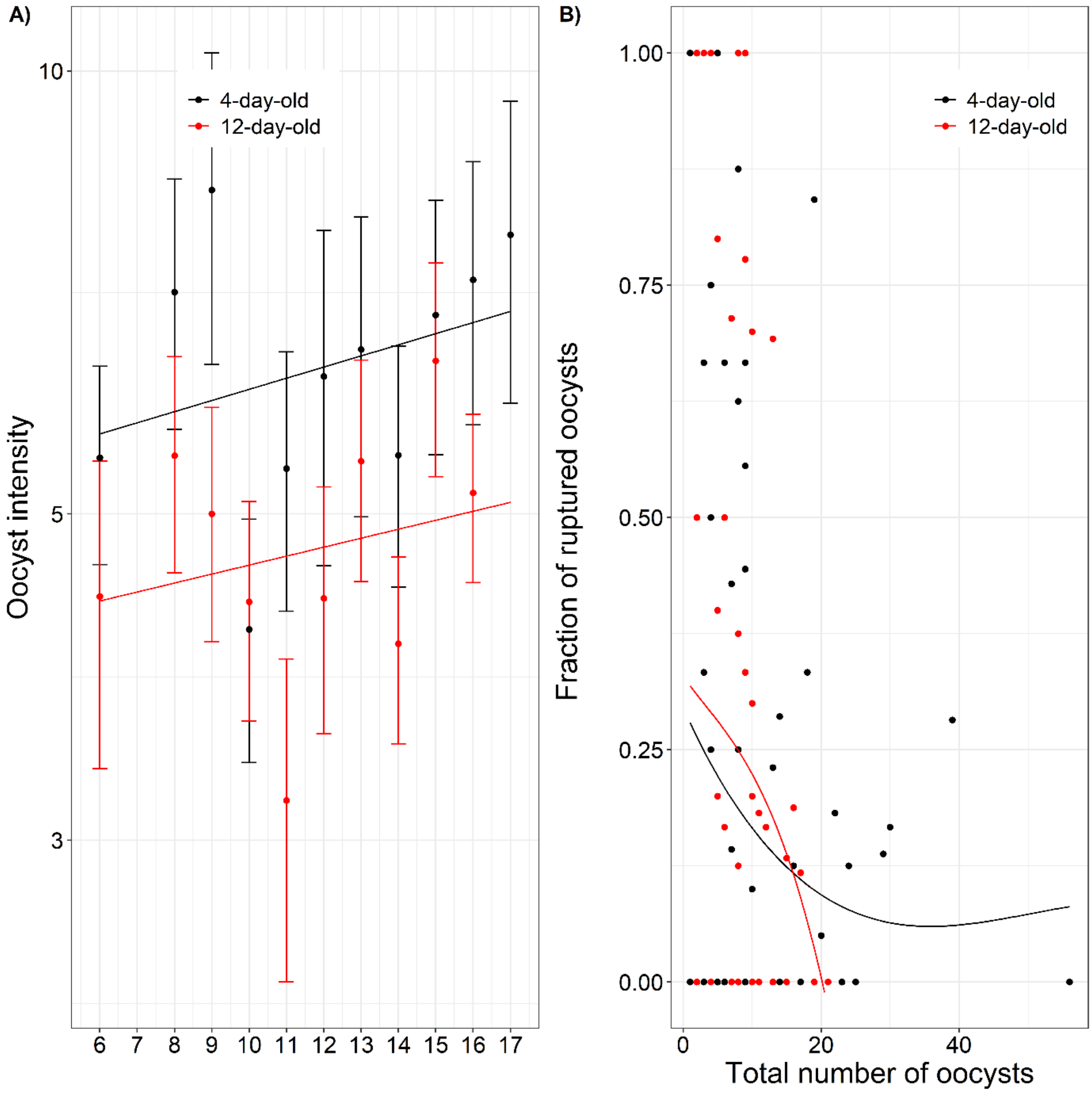


**Figure S5:** (A) Oocyst intensity as a function of time post-infection in days for each of the two age class. (B) The fraction of ruptured oocysts as a function of the total number of oocysts in individual mosquitoes for each of the two age class. The fraction of ruptured oocysts is the number of ruptured oocysts out of the total number of oocysts in the midguts of infected mosquitoes from 6-17 dpbm. Each black dot represent a mosquito individual and the lines represents the best-fit relationship.
